# Expression dynamics and functional characterization of the *pigmented and non-diapausing egg* gene (*pnd*) and *pnd-2*, responsible for the initiation of embryonic diapause in the silkworm, *Bombyx mori*

**DOI:** 10.1101/2025.08.17.667111

**Authors:** Hayato Yamada, Shouhei Kihara, Teruyuki Niimi, Keiko Kadono-Okuda, Toshinobu Yaginuma

## Abstract

Diapause in insects is a developmental arrest that serves as an adaptation to severe environmental conditions. In the silkworm *Bombyx mori*, embryonic diapause in the next generation is maternally induced by the diapause hormone (DH). However, the downstream factors responsible for initiating diapause have remained unidentified. The *pnd* and *pnd-2* mutants produce *p*igmented and *n*on-*d*iapausing eggs despite intact DH signaling, indicating that the *pnd* and *pnd-2* genes act downstream as potential initiators of diapause. Here, we analyzed their expression dynamics and functions. Expression analyses showed that *pnd* was strongly upregulated in diapause-destined eggs in a DH-signal-dependent manner and was suppressed by HCl treatment that prevents diapause initiation. In contrast, *pnd-2* exhibited a transient expression peak independent of DH-signal and was unaffected by HCl treatment. RNAi-mediated knockdown of either gene inhibited the diapause initiation, and injection of *pnd* mRNA into non-diapause-destined eggs induced a diapause-like state in embryos. These findings demonstrate that *pnd* and *pnd-2* are required downstream of the DH-signal to initiate diapause, providing key insights into the molecular mechanism underlying embryonic diapause induction in *B. mori*.

**Graphical Abstract:** 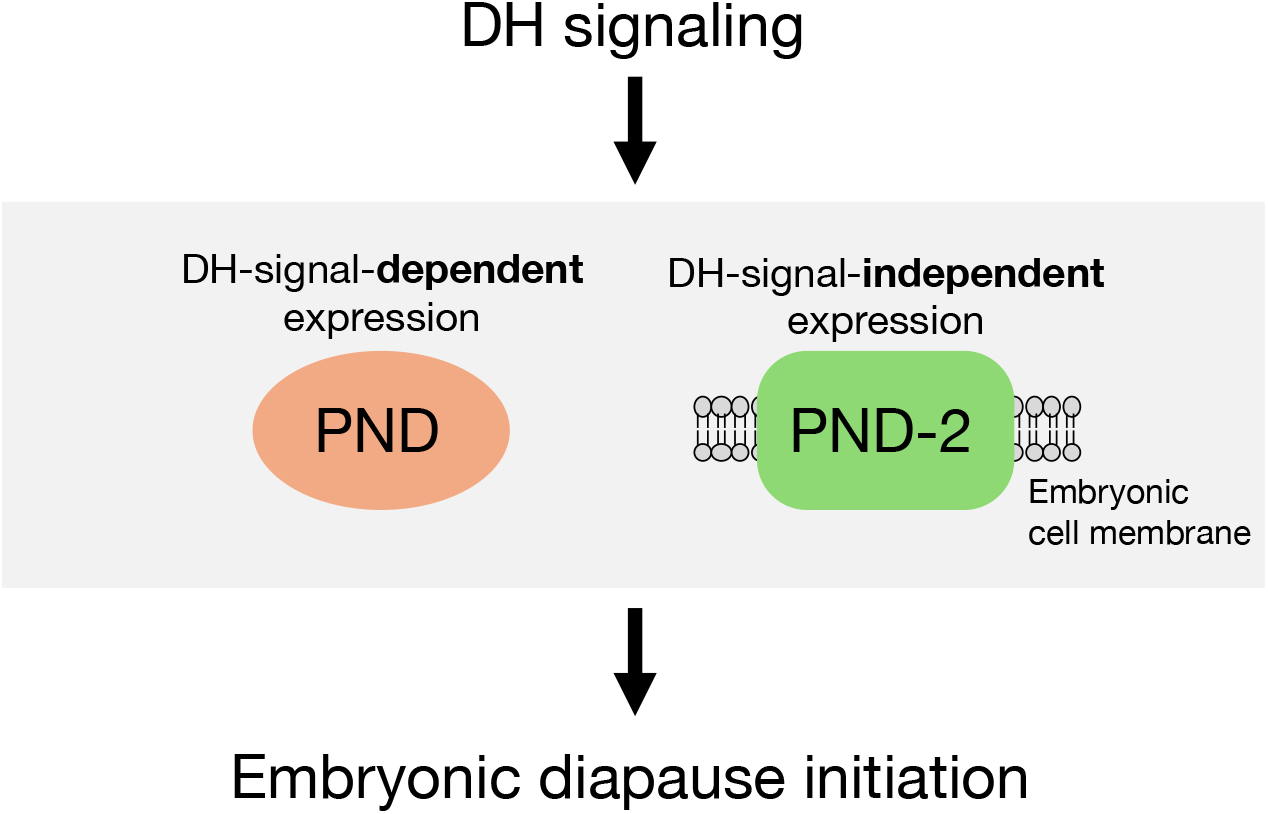

## Introduction

Insects have evolved a developmental arrest known as diapause to survive unfavorable conditions such as winter (Denlinger, 2022; Koštál et al., 2017). This adaptive state prolongs survival by suppressing metabolic activity and development, and its molecular mechanisms have attracted interest in both basic and applied research (Takahashi et al., 2020).

In the silkworm *Bombyx mori*, diapause occurs during early embryogenesis, immediately after mesoderm segregation and telson differentiation (2–3 days after oviposition under 25^°^C), when embryonic cells arrest in the G2 phase (Miya, 2003; Nakagaki et al., 1991; Yamashita and Yaginuma, 1991). Embryonic diapause is determined by maternal diapause hormone (DH) signaling (Shiomi et al., 2015; Yamashita and Hasegawa, 1985). DH secreted from the suboesophageal ganglion of female pupae in the mid-pupal stage enters the hemolymph and binds to the diapause hormone receptor (DHR) on developing oocytes; eggs whose oocytes receive DH enter diapause after oviposition, whereas those without DH input complete embryogenesis within approximately 10 days and then larvae hatch (Hasegawa, 1957; Homma et al., 2006; Yamashita, 1996). *B. mori* races are classified as univoltine, bivoltine, or multivoltine: univoltine races consistently produce diapause eggs, multivoltine races produce non-diapause eggs, and in bivoltine races maternal temperature and photoperiod during the mother’s embryonic stage regulate DH release, yielding either diapause or non-diapause eggs (Yamashita and Hasegawa, 1985).

In addition to inducing embryonic diapause, DH action can also be monitored by serosal pigmentation, with diapause eggs turning from pale yellow to dark brown within 2–3 days after oviposition (Sonobe and Ohnishi, 1970; Yamashita and Hasegawa, 1964). The maternal endocrine pathway that links environmental cues such as temperature and photoperiod to DH secretion acting on developing oocytes has been partly elucidated (Sato et al., 2014; Shiomi et al., 2015; Tsuchiya et al., 2020). However, the molecular mechanisms acting downstream of DH signaling and initiating embryonic diapause remain largely unknown. Notably, two mutant races were discovered in which eggs show normal serosal pigmentation, yet continue embryonic development and hatch without entering diapause; these are commonly referred to as “pigmented and non-diapausing eggs” (Katsumata, 1968; Kihara, 1979). Positional cloning identified the responsible genes for these mutants as *pnd* (*pigmented and non-diapausing egg*) and *pnd-2* (Kidokoro et al., 2010; K. Kadono-Okuda, unpublished results). Because serosal pigmentation occurs normally in *pnd* and *pnd-2* mutants, upstream DH signaling is intact. These observations suggest that *pnd* and *pnd-2* act downstream of DH signaling as effectors initiating diapause.

In this study, we examined the mRNA expression dynamics of *pnd* and *pnd-2* during embryogenesis and performed both loss- and gain-of-function analyses to verify their roles, demonstrating that these genes act as positive regulators downstream of DH signaling and drive the initiation of embryonic diapause in *B. mori*.

## Results

### Cloning and structural analysis of *pnd* and *pnd-2*

To analyze the function of *pnd* and *pnd-2*, we re-cloned their open reading frames, *pnd* (387 bp) and *pnd-2* (1530 bp), by RT-PCR using cDNA prepared from diapause-destined eggs of the bivoltine race Daizo. Both sequences were identical to previously deposited sequences derived from the p50T silkworm race (GenBank accession nos. LC859351 and LC860026), confirming their conservation across genetic backgrounds.

The predicted amino acid sequences indicate that PND is an approximately 14-kDa protein with an N-terminal signal peptide (residues 1–22), whereas PND-2 is an approximately 56-kDa protein with an N-terminal signal peptide (residues 1–16) and a single transmembrane domain (residues 254–276) (Fig. 1A). These structural characteristics suggest that PND functions as a secreted ligand-like protein, and PND-2 as a membrane-associated receptor-like protein.

**Fig. 1.**
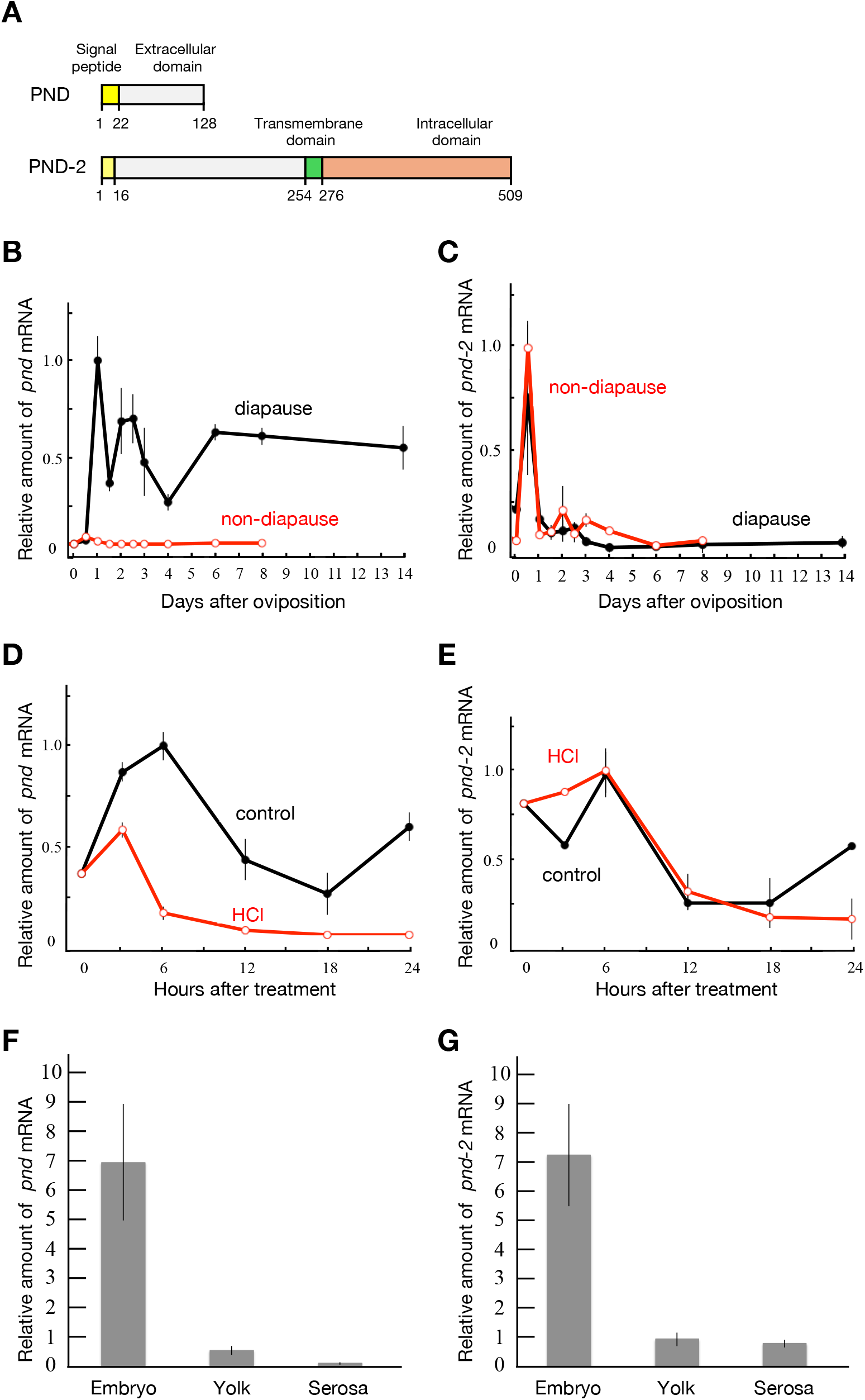
Domain architecture and expression profiles of *pnd* and *pnd-2*. (A) Schematic representation of predicted domain architecture of PND and PND-2. Signal peptides (yellow), extracellular domains (gray), a transmembrane domain (green), and an intracellular domain (orange) are indicated. Amino acid positions are shown. (B, C) Post-oviposition expression profiles of *pnd* and *pnd-2* mRNA in diapause-destined and non-diapause-destined eggs (Daizo race), quantified by qRT-PCR at the indicated days after oviposition. In non-diapause eggs, larvae hatched at 9–10 days after oviposition. Expression levels were normalized to the internal control gene *rp*^*49*^ using the ΔΔCt method. (D, E) Expression profiles of *pnd* and *pnd-2* mRNA after HCl or water (control) treatment in diapause-destined eggs. Eggs were treated at 24 h post-oviposition and sampled at the indicated time points after treatment (defined as 0 h). Expression levels were quantified by qRT-PCR and normalized to *rp49*. (F, G) Tissue expression profiles of *pnd* and *pnd-2* mRNA in embryo, yolk, and serosa dissected from diapause-destined eggs (Daizo race) at 5–6 days after oviposition. Expression levels were quantified by qRT-PCR and normalized to those in whole eggs (not shown). Data are means ± SEM; *n* = 2 for (B–E), *n* = 3 for (F, G).

### Expression dynamics of *pnd* and *pnd-2*

To characterize the temporal expression profiles of *pnd* and *pnd-2*, we quantified their mRNA levels in diapause- and non-diapause-destined eggs at multiple time points after oviposition by qRT-PCR (Fig. 1B, C). For *pnd*, mRNA expression was low at 0 and 12 h post-oviposition in both diapause and non-diapause eggs. In diapause eggs, *pnd* expression increased markedly at 24 h and remained consistently high through day 14. In contrast, its levels in non-diapause eggs remained low until day 8, and larvae hatched at days 9–10. For *pnd-2*, mRNA expression was low at 0 h in both diapause and non-diapause eggs. It exhibited a transient increase at 12 h, and then returned to low levels thereafter, remaining low at all subsequent observed time points for both diapause eggs and non-diapause eggs. These results suggest that *pnd* is expressed in response to DH signaling, whereas *pnd-2* is transiently expressed in a developmentally programmed manner.

Embryonic diapause initiation in silkworm eggs can be averted by HCl treatment between 20 and 24 h after oviposition (Tsurumaru, 2010; Yamashita and Yaginuma, 1991). To examine how HCl treatment affects *pnd* and *pnd-2* mRNA expression, diapause-destined eggs were treated with HCl or tap water (control) at 24 h after oviposition and sampled at multiple time points over the following 24 h. Their mRNA expression was then analyzed by qRT-PCR. *pnd* mRNA expression markedly decreased within 6 h after HCl treatment and remained low, whereas expression in control eggs remained high throughout the 24 h observation period (Fig. 1D). For *pnd-2*, mRNA expression gradually decreased during the 24 h observation period in both HCl-treated and control eggs, with no substantial difference between treatments (Fig. 1E). These results indicate that *pnd*, but not *pnd-2*, is transcriptionally downregulated in response to HCl treatment-induced diapause-averting conditions.

### Tissue distribution of *pnd* and *pnd-2*

We next examined where within the eggs *pnd* and *pnd-2* mRNA accumulate. We quantified their mRNA in the embryo, yolk, and serosa dissected from diapause-destined eggs at 5–6 days post-oviposition by qRT-PCR (Fig. 1F, G). Both genes exhibited predominant expression in the embryo, with little or no detectable expression in the yolk and serosa. This distribution suggests that *pnd* and *pnd-2* act mainly within the embryo to regulate diapause initiation.

### Loss-of-function analysis of *pnd* and *pnd-2*

We performed embryonic RNAi in diapause-destined eggs by injecting dsRNA targeting *pnd* or *pnd-2* within 5 h after oviposition, with ultrapure water injection used as a control (Fig. 2). Twenty days after injection, eggs were fixed in Carnoy’s solution. After chorions were removed, embryos were stained with carbolic-thionine solution to determine developmental stage and whether embryos had progressed beyond the diapause stage. All control embryos remained at the diapause stage (0/50 developed beyond diapause). In contrast, 16.0% (4/25) of *pnd* dsRNA-injected embryos and 56.5% (26/46) of *pnd-2* dsRNA-injected embryos developed beyond the diapause stage. These results confirm Kidokoro et al. (2010) and show that knockdown of *pnd* or *pnd-2* inhibits diapause initiation, leading to continued embryonic development as in non-diapause eggs.

**Fig. 2.**
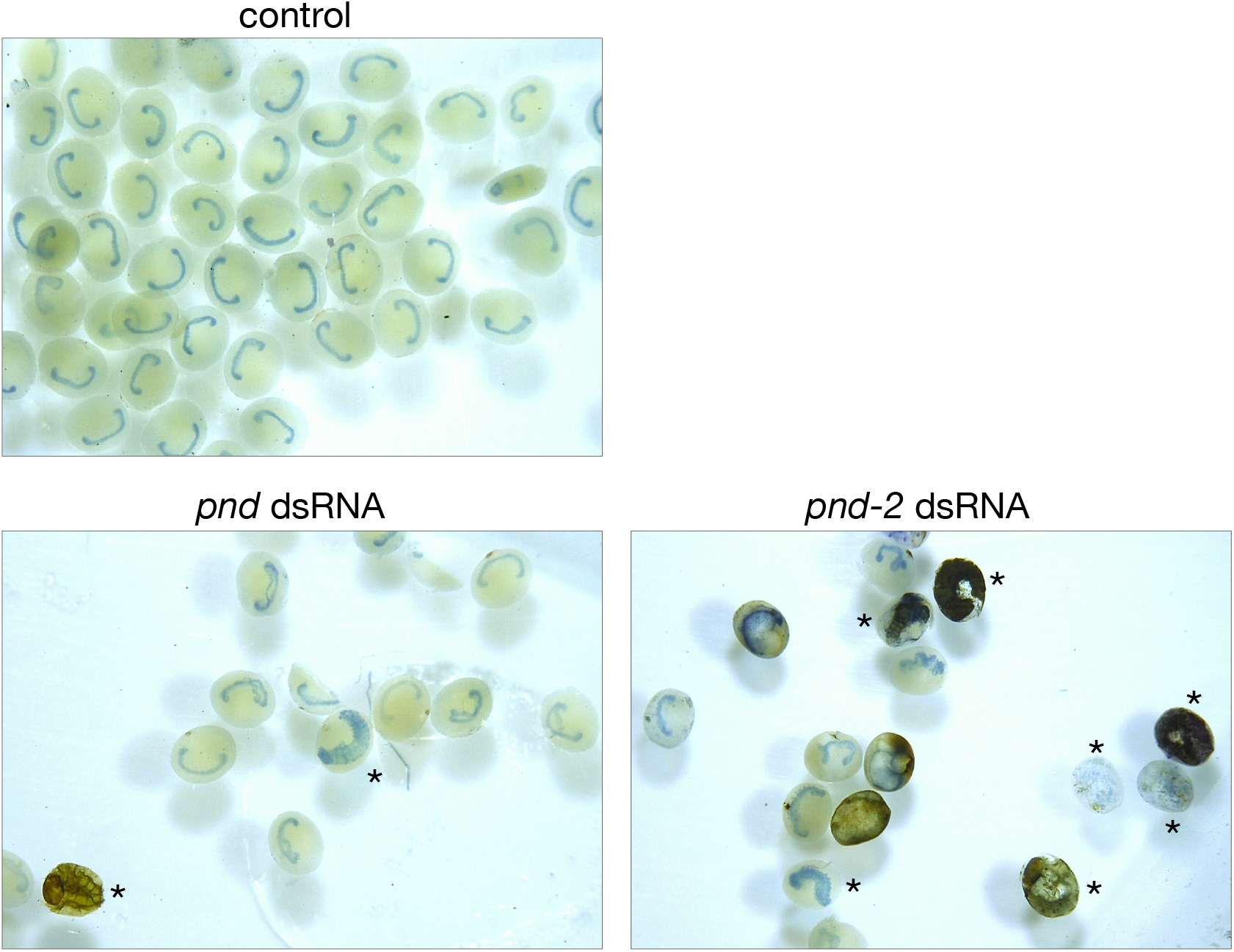
Embryonic RNAi analysis of *pnd* and *pnd-2*. Representative images of diapause-destined eggs (0081w-1) injected with ultrapure water (control; top), *pnd* dsRNA (bottom left), or *pnd-2* dsRNA (bottom right). Eggs were fixed 20 days after injection, dechorionated, stained with carbolic-thionine, and imaged. Asterisks indicate embryos that developed beyond the diapause stage.

### Gain-of-function analysis of *pnd*

Loss-of-function analyses indicate that both *pnd* and *pnd-2* are required for the initiation of embryonic diapause, and the expression analyses showed that high *pnd* expression—suppressed under diapause-averting HCl treatment—is specifically associated with diapause induction. Here, we conducted a gain-of-function experiment by injecting synthesized *pnd* mRNA into eggs of multivoltine, non-diapause race N_4_ within 4 h after oviposition, with ultrapure water injection used as a control (Fig. 3). 12 days after injection, embryonic morphology was checked for describing their developmental stage. All control embryos developed to the pre-hatching or hatching stage (0/37 remained at the diapause stage). In contrast, 77.8% (28/36) of *pnd* mRNA-injected eggs exhibited diapause-like arrest characteristic of diapause-stage embryos. These results demonstrate that exogenous *pnd* expression is sufficient to induce a diapause-like state in otherwise non-diapause embryos, supporting its role as a key driver of diapause initiation.

**Fig. 3.**
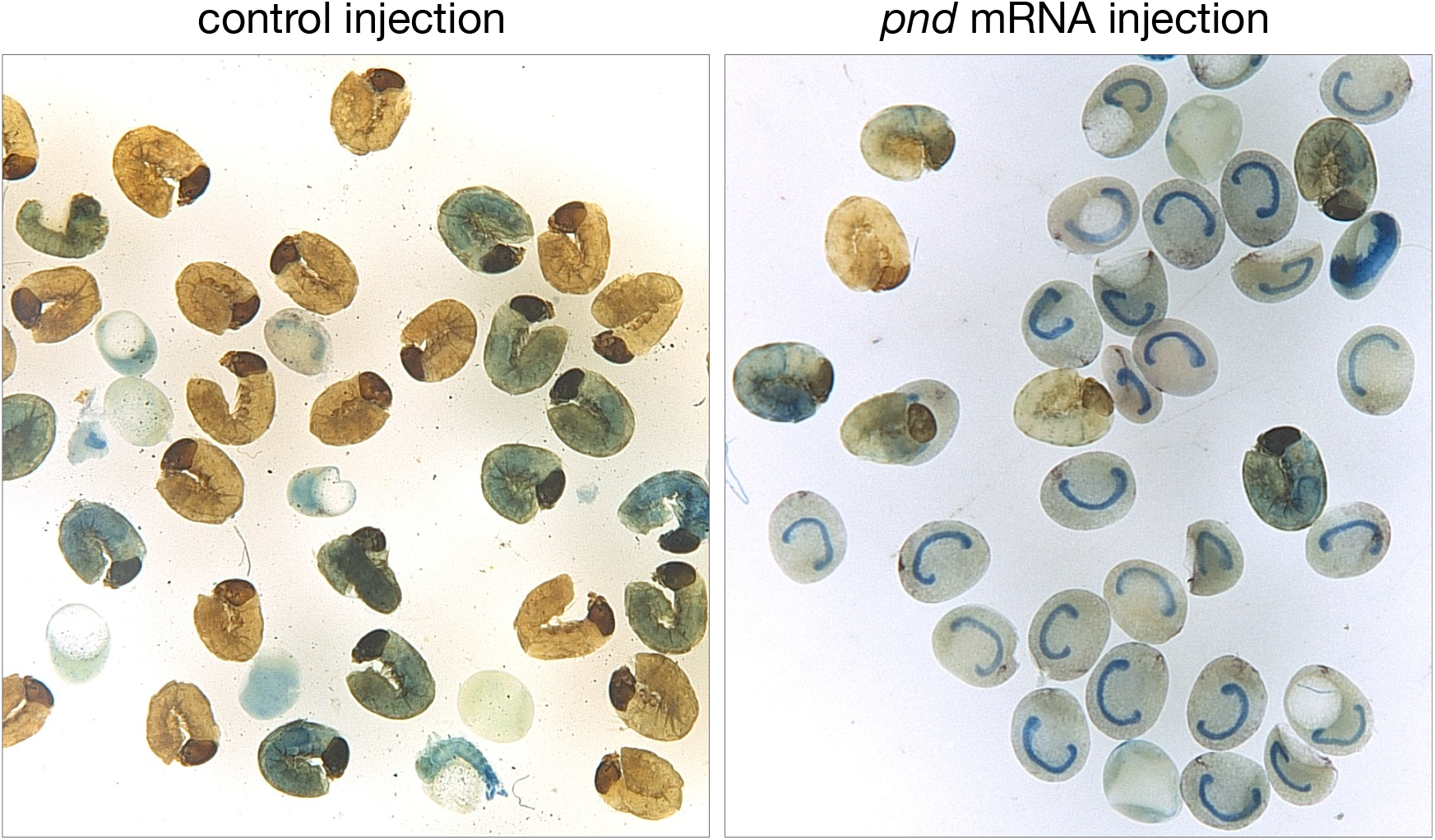
Effects of *pnd* mRNA injection on embryonic development in eggs from a non-diapause race. Representative images of eggs from a non-diapause race N_4_ injected with ultrapure water (control; left) or synthesized *pnd* mRNA (right). Eggs were fixed 12 days after injection, dechorionated, stained with carbolic-thionine, and examined for embryonic morphology.

## Discussion

In this study, we examined the roles of *pnd* and *pnd-2* in embryonic diapause initiation in *B. mori* through post-oviposition expression profiling, loss-of-function experiments using embryonic RNAi, and gain-of-function experiments by *pnd* mRNA microinjection. Our results demonstrated that both genes acted downstream of DH-signal as positive regulators of diapause initiation. Expression analyses revealed that *pnd* was transcriptionally induced by the DH-signal, whereas *pnd-2* exhibited a transient, developmentally programmed expression peak independent of the DH-signal. Suppression of *pnd* expression under diapause-averting HCl treatment further supports that high *pnd* expression is required for diapause initiation. RNAi knockdown of either gene inhibited diapause initiation, with a stronger effect for *pnd-2*, likely due to their distinct molecular roles: PND is a secreted, ligand-like protein that can act across cells, whereas PND-2 is a membrane-associated, receptor-like protein that must be present in each cell. These functional distinctions are consistent with mosaic embryo analyses of *pnd* and *pnd-2* mutants (Kobayashi, 1991): in mosaics containing wild-type and *pnd* cells, entire embryos remained at the diapause stage; in mosaics containing wild-type and *pnd-2* cells, some cells remained at the diapause stage whereas others developed beyond the diapause stage, resulting in embryonic lethality. Microinjection of *pnd* mRNA efficiently induced a diapause-like state in embryos of a non-diapause race, further supporting its role as a secreted factor.

Together, these findings support a model in which the secreted ligand PND and the membrane-associated receptor PND-2 interact directly or indirectly within embryonic tissues after oviposition, where both genes are predominantly expressed, to initiate the diapause cascade. Future work should address two key questions. First, how does DH signaling regulate *pnd* expression after oviposition? Second, how do PND–PND-2 interaction and downstream pathways drive diapause initiation?

## Materials & methods

### Silkworm races and maintenance conditions

The following *Bombyx mori* races were used in this study: Daizo (a bivoltine race maintained at Nagoya University), Kinshu × Showa (a commercial hybrid race producing diapause eggs), the mutant race 0081w-1 (a diapause race producing eggs without serosal pigmentation, obtained from the National Bio-Resource Project [NBRP] at Kyushu University), and N_4_ (a multivoltine, non-diapause race maintained at Nagoya University). To induce diapause or non-diapause eggs in the bivoltine race, the parental generation was incubated during its embryonic stage under 24LL at 25^°^C or 24DD at 15^°^C, respectively. Except for these incubation conditions used to induce diapause or non-diapause eggs, all eggs were incubated at 25^°^C, and sampling was performed at appropriate time points relative to oviposition. Silkworm larvae were reared on mulberry leaves at 25^°^C under a 12L:12D photoperiod.

### Cloning and domain analysis

#### cDNA preparation

Total RNA was extracted from diapause-destined eggs (2 days after oviposition) of the biovoltine race Daizo using TRIzol Reagent (Gibco BRL). Approximately 50–100 mg of eggs (one egg equivalent to about 0.5 mg) were homogenized, and RNA was purified by chloroform extraction, isopropanol precipitation, and ethanol washing. The RNA was treated with DNase I, followed by an additional purification step using TRIzol, and finally dissolved in ultrapure water. First-strand cDNA was synthesized from the purified total RNA using SuperScript II (Invitrogen).

#### PCR amplification and cloning

Full-length *pnd* and *pnd-2* cDNA were amplified by PCR using gene-specific primer pairs pnd_F/pnd_R and pnd2_F/pnd2_R, respectively, with the above cDNA as a template (Table 1). PCR was performed using KOD-Plus-Neo (TOYOBO) with the following touchdown program: an initial denaturation at 94^°^C for 2 min; 15 cycles of 98^°^C for 10 s, annealing from 74^°^C to 58^°^C with a 4^°^C decrement every three 3 cycles for 30 s, and 68^°^C for 90 s; followed by 50 cycles of 98^°^C for 10 s, 52^°^C for 30 s, and 68^°^C for 90 s; and a final extension at 68^°^C for 7 min. PCR products were confirmed and purified by agarose gel electrophoresis and subjected to A-tailing using 10 × A-attachment mix (TOYOBO). The *pnd* fragment was cloned into the EcoRV site of the pBluescript KS(+) vector (Stratagene), and the *pnd-2* fragment was cloned into the pTA2 vector (TOYOBO). Recombinant plasmids were transformed into XL-1 Blue competent cells (Stratagene) and selected on LB agar plates containing ampicillin. Plasmid DNA was purified from positive colonies using the Axyprep Plasmid Miniprep Kit (Axygen).

**Table 1.**
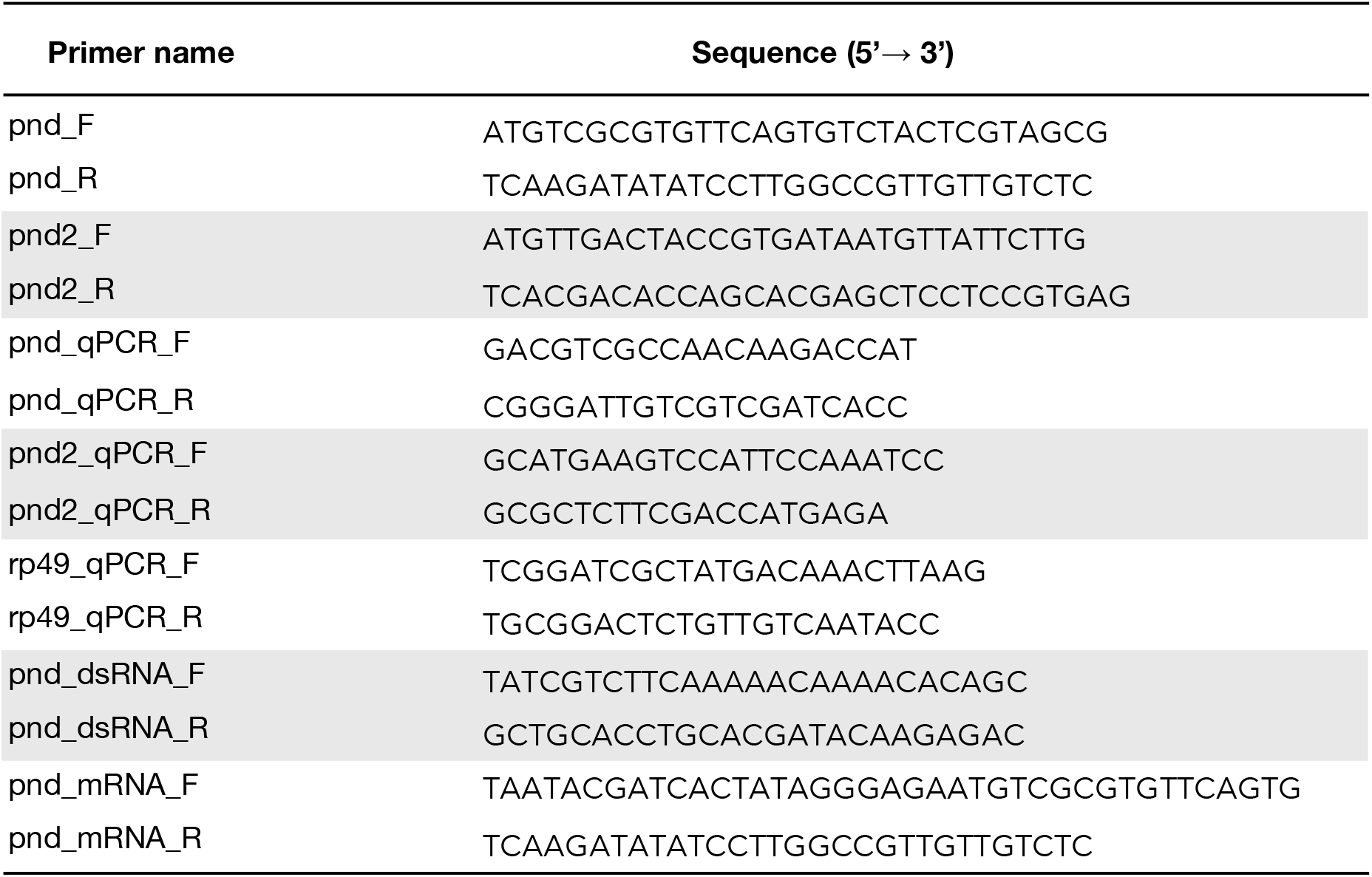
Oligonucleotide primers used in this study.

#### Sequence determination and domain analysis

Plasmid DNA was sequenced using the BigDye Terminator v3.1 Cycle Sequencing kit (Applied Biosystems) on an ABI PRISM 3100 Avant Genetic Analyzer (Applied Biosystems). The obtained nucleotide sequences were verified using GENETYX (GENETYX Corporation). Predicted amino acid sequences of PND and PND-2 were further analyzed for domain prediction using InterProScan (Paysan-Lafosse et al. 2023, https://www.ebi.ac.uk/interpro/).

### mRNA expression analysis

#### Egg preparation

Eggs of the bivoltine race Daizo were used to analyze *pnd* and *pnd-2* mRNA expression in diapause and non-diapause eggs. Diapause eggs were sampled from 0 to 14 days after oviposition, and non-diapause eggs were sampled from 0 to 9 days after oviposition.

Eggs of the bivoltine hybrid commercial race (Kinshu × Showa) were used to analyze *pnd* and *pnd-2* mRNA expression in diapause eggs in which diapause initiation was prevented by HCl treatment. Diapause-destined eggs were treated 24 h after oviposition with HCl (specific gravity 1.110) at 25^°^C for 90 min under room temperature, or with tap water as a control, and sampled at 0, 3, 6, 12, 18, and 24 h after treatment.

Diapause eggs of Daizo were used to measure *pnd* and *pnd-2* mRNA levels in the embryos, yolk cells, and serosa. These tissues were dissected from 20–30 eggs at 5 and 6 days after oviposition, when these tissues were distinguishable and could be separated under a stereomicroscope using forceps in phosphate-buffered saline (PBS). Each tissue was immediately frozen in liquid nitrogen and stored at -80^°^C. Whole eggs were collected in parallel with individual tissues and used as a reference sample for relative quantification in subsequent quantitative RT-PCR (qRT-PCR) analyses.

#### Quantitative RT-PCR (qRT-PCR)

Total RNA was extracted, and first-strand cDNA was synthesized as described above. This cDNA was used as a template for qRT-PCR, which was conducted using a StepOnePlus Real-Time PCR System (Applied Biosystems) with SYBR Select Master Mix (Applied Biosystems). Primer pairs pnd_qPCR_F/pnd_qPCR_R, pnd2_qPCR_F/pnd2_qPCR_R, and rp49_qPCR_F/rp49_qPCR_R were used to amplify *pnd, pnd-2*, and the ribosomal protein gene *rp49*, respectively (Table 1). For analyses across time points after oviposition and after HCl treatment, *rp49* was used as an internal control, and *pnd* and *pnd-2* mRNA levels were normalized using the ΔΔCt method. For analyses across tissues, where *rp49* expression levels may vary among tissues, relative quantification was performed by normalizing to the expression level in whole eggs.

## Embryonic RNA interference (RNAi)

Total RNA was extracted from eggs of the bivoltine silkworm race p50T, collected 24–96 h post-oviposition, and reverse-transcribed to synthesize first-strand cDNA. This cDNA served as the template for PCR amplification of *pnd* and *pnd-2* fragments for dsRNA synthesis. For *pnd*, the primer pair pnd_dsRNA_F/pnd_dsRNA_R was used (Table 1); for *pnd-2*, a primer pair was used whose sequences are available upon request from co-author K. K.-O. The amplified fragments were cloned into the pSTBlue-1 vector (Novagen), and the resulting plasmids were PCR-amplified using T7 universal primers to generate DNA fragments flanked by T7 promoter sequences. These fragments served as templates for *in vitro* transcription using the MEGAscript T7 kit (Ambion), and the sense and antisense RNA strands were annealed by heat denaturation and cooling to yield double-stranded RNA (dsRNA).

Each dsRNA targeting *pnd* or *pnd-2* was prepared at 2 µg/µL and injected separately into diapause-destined eggs of the mutant race 0081w-1, which lacks ommochrome pigmentation in the serosal cells due to a defect in kynurenine 3-hydroxylase, an enzyme in the ommochrome biosynthesis pathway. Injections were performed within 5 h post-oviposition, following the microinjection protocol described by Yamada et al. (2025). Ultrapure water was injected as a control.

### mRNA microinjection

DNA fragments for *in vitro* transcription were PCR-amplified using the primer pair pnd_mRNA_F/pnd_mRNA_R, which contained T7 promoter sequences, and KOD-FX-Neo (TOYOBO), with the pBluescript KS(+) plasmid carrying the full-length *pnd* sequence serving as the template (Table 1). The PCR products were confirmed and purified by agarose gel electrophoresis and then used as templates to synthesize *pnd* mRNA with the mMESSAGE mMACHINE T7 Ultra Kit (Life Technologies).

The synthesized *pnd* mRNA was prepared at 1.4 µg/µL and injected into eggs of the multivoltine, non-diapause race N_4_ within 4 h post-oviposition. Microinjection procedures were identical to those used for the RNAi experiments. Ultrapure water was injected as a control.

### Assessment of embryonic development and diapause traits in injected embryos

Eggs injected with *pnd* or *pnd-2* dsRNA were incubated at 25^°^C and fixed 20 days after injection, whereas those injected with synthesized *pnd* mRNA were fixed 12 days after injection. Fixation was performed overnight in Carnoy’s solution, a mixture of ethanol, chloroform, and acetic acid in a 6:3:1 ratio by volume. The chorions were removed after fixation, and the embryos were stained with carbolic-thionine diluted in 80% ethanol to visualize the embryos and then made transparent with benzene. The embryos were examined under a stereomicroscope (VHX-900, KEYENCE) to assess morphological changes related to developmental stage or diapause stage (Yaginuma et al., 1990).

## Data availability

The nucleotide sequences of *pnd* and *pnd-2* have been deposited in GenBank under accession nos. LC859351 and LC860026, respectively. Further data are available from the corresponding author upon reasonable request.

## Acknowledgments

We thank Prof. M. Ikeda and late Emeritus Prof. M. Kobayashi at Nagoya University, Emeritus Prof. O. Yamashita at Nagoya University and Chubu University, and late Emeritus Prof. M. Kobayashi at the University of Tokyo for their encouragement during this study. Silkworm races used in this study were assisted in part by the NBRP of the MEXT, Japan.

## Author contributions

**Hayato Yamada:** Conceptualization, Writing – original draft, Visualization. **Shouhei Kihara:** Investigation, Data curation. **Teruyuki Niimi:** Methodology, Investigation, Conceptualization. **Keiko Kadono-Okuda:** Writing – review & editing. **Toshinobu Yaginuma:** Supervision, Project administration, Funding acquisition, Conceptualization.

## Funding

This work was supported partially by a JSPS (Japan Society for the Promotion of Science) KAKENHI grant (numbers 21248007 and 24380033) to T. Y.

## Declaration of competing interest

The authors declare no competing interests.

## Notes

### Competing Interest Statement

The authors have declared no competing interest.

